# Transplantation of high fat fed mouse microbiota into zebrafish larvae identifies MyD88-dependent acceleration of hyperlipidaemia by Gram positive cell wall components

**DOI:** 10.1101/2021.04.28.441900

**Authors:** Pradeep Manuneedhi Cholan, Simone Morris, Kaiming Luo, Jinbiao Chen, Jade A Boland, Geoff W McCaughan, Warwick J Britton, Stefan H Oehlers

## Abstract

Gut dysbiosis is an important modifier of pathologies including cardiovascular disease but our understanding of the role of individual microbes is limited. Here, we have used transplantation of mouse microbiota into microbiota-deficient zebrafish larvae to study the interaction between members of a mammalian high fat diet-associated gut microbiota with a lipid rich diet challenge in a tractable model species. We find zebrafish larvae are more susceptible to hyperlipidaemia when exposed to the mouse high fat-diet-associated microbiota and that this effect can be driven by two individual bacterial species fractionated from the mouse high fat-diet-associated microbiota. We find *Stenotrophomonas maltophilia* increases the hyperlipidaemic potential of chicken egg yolk to zebrafish larvae independent of direct interaction between *S. maltophilia* and the zebrafish host. Colonisation by live, or exposure to heat-killed, *Enterococcus faecalis* accelerates hyperlipidaemia via host Myd88 signalling. The hyperlipidaemic effect is replicated by exposure to the Gram positive Toll-like receptor agonists peptidoglycan and lipoteichoic acid in a MyD88-dependent manner. In this work, we demonstrate the applicability of zebrafish as a tractable host for the identification of gut microbes that can induce conditional host phenotypes via microbiota transplantation and subsequent challenge with a high fat diet.

## Introduction

The metagenome encoded by the gut microbiome is an essential component of the animal digestive system [1, 2]. Microbes can affect digestion directly through the breakdown of indigestible material such as fibre to short chain fatty acids and indirectly by stimulating the differentiation of the intestinal epithelium.

The microbiome is shaped by host and environment selective pressures [3, 4]. Lipid-rich Western diets that cause obesity in susceptible individuals are associated with gut dysbiosis which can drive an unwanted increase in nutrient absorption and epithelial leakiness [5]. Transplantation of human lean and obese microbiota into germ-free mice transmits these phenotypes across host mammalian species [6]. While the germ-free mouse has been invaluable for the study of host-microbe interactions through gnotobiotic and transplantation studies, there is a need for faster and more accessible models that serve as functional screening tools for disease-associated microbiota.

Zebrafish embryos require microbial colonisation during development for full physiological digestive function and are a simple platform for studying host-microbiota-environment interactions [7, 8]. Zebrafish embryos develop *ex utero* within a chorion that can be surface sterilised and are tolerant of antibiotics in their media making them a technically simple model for raising germ-free experimental subjects. Probiotic transfer has been shown to increase resistance to pathogen colonisation [9]. The zebrafish gut microbiome is sensitive to challenge with a HFD, feeding larval zebrafish a diet supplemented with 10% fat for 25 days altered the distribution of bacterial phyla in the gut which correlated with increased expression of host inflammatory genes compared to larvae fed the control diet [3].

Microbiota transplantations have been previously performed from mice and humans to zebrafish demonstrating the feasibility of using the zebrafish as an in vivo screening tool for studying the effects of defined microbiota on host physiology [10]. Here we use this technically simple model to rapidly identify specific members of the mouse HFD-associated microbiota that accelerate a diet-induced hyperlipidaemic phenotype in zebrafish embryos and the mechanisms by which these pathobionts interact with the zebrafish host.

## Results

### Transplantation of microbiota from HFD fed mice accelerates hyperlipidaemia in zebrafish embryos

Microbiome-depleted (MD) zebrafish embryos were exposed to mouse faecal microbiota preparations generated from mice fed a conventional chow diet or HFD from 3-5 dpf (days post fertilization) and challenged with chicken egg yolk feeding from 5-7 dpf (Figure 1A).

**Figure 1:**
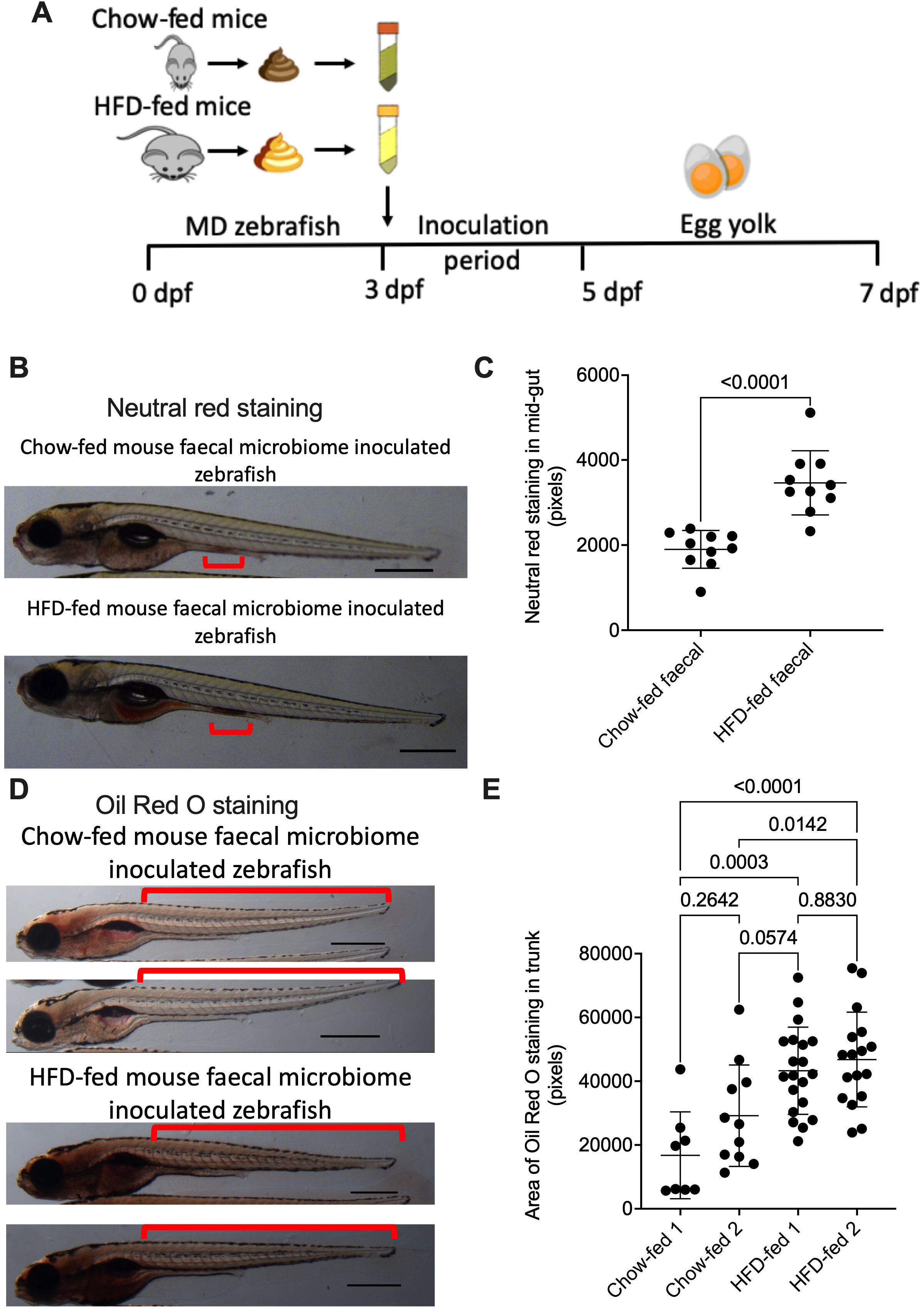
Transplantation of microbiota from HFD-fed mice accelerates hyperlipidaemia in zebrafish embryos. (a) Schematic describing method of faecal microbiome inoculation in MD zebrafish and chicken egg yolk challenge diet. (b) Representative images of neutral red staining in mid-gut of 5 dpf zebrafish embryos. Red brackets indicate mid-gut region used for quantification. (c) Quantification of neutral red staining area in mid-gut of 5 dpf zebrafish embryos. (d) Representative images of Oil Red O staining of 7 dpf chow-fed and HFD-fed mouse faecal microbiome-inoculated zebrafish embryos. Red brackets indicate tail region posterior to the swim bladder used for quantification (e) Quantification of trunk vascular Oil Red O staining from the tail region posterior to the swim bladder in MD zebrafish inoculated with mouse faecal microbiota and challenged with chicken egg yolk diet from 5-7 dpf. Scale bar represents 500 μm. Results are expressed as mean ± SD.

We first investigated if colonisation with the control or HFD-fed mouse microbiota affected zebrafish intestinal physiology at 5 dpf before the onset of exogenous feeding. The absorptive activity of zebrafish midgut lysosome-rich intestinal epithelial cells can be visualised by neutral red staining, previous studies have demonstrated that this phenotype requires microbial colonisation and is impeded by inflammation [11, 12]. We used neutral red staining visually to examine the absorptive function of colonised embryos and observed higher staining in 5 dpf larvae colonised with the HFD-fed microbiota compared to chow diet colonised controls (Figure 1B and 1C).

We next collected colonised zebrafish larvae and stained with Oil Red O to visualise neutral lipids but did not observe any differences in lipid staining. To determine if colonisation with the HFD-fed mouse microbiota affected the zebrafish response to challenge with a chicken egg yolk challenge [13], we challenged 5 dpf zebrafish with chicken egg yolk and stained with Oil Red O to visualise neutral lipids (Figure 1D). Zebrafish larvae colonised with the faecal microbiota from HFD fed mice had more vascular Oil Red O staining after 2 day of chicken egg yolk feeding challenge compared to zebrafish larvae colonised with the faecal microbiota from chow diet fed mice (Figure 1E).

These data demonstrate the responsiveness of MD zebrafish larvae to the transferable effects of dysbiotic mammalian gut microbiome-associated microbiota when challenged with a complex environmental stimulus in the form of lipid-rich feeding.

### Identification of individual microbes with pathobiont activity

To isolate individual species that would be amenable to *in vitro* handling and growth, faecal homogenate supernatants were plated on LB agar and grown at 28°C to select for species that would be easily handled and most likely to colonise zebrafish embryos (Figure 2A). The best growing isolate in LB broth culture from two chow diet and HFD fed faecal preparations were selected and sequenced. We identified *Enterococcus faecalis* strain YN771 (*E.f* on figures) and *Stenotrophomonas maltophilia* strain CD103 (*S.m*) from HFD-fed mouse faecal lysate, and *Escherichia* species PYCC8248 (E.s) and *Escherichia coli* strain Y15-3 (*E.c*) from chow diet-fed mouse faecal lysate.

**Figure 2:**
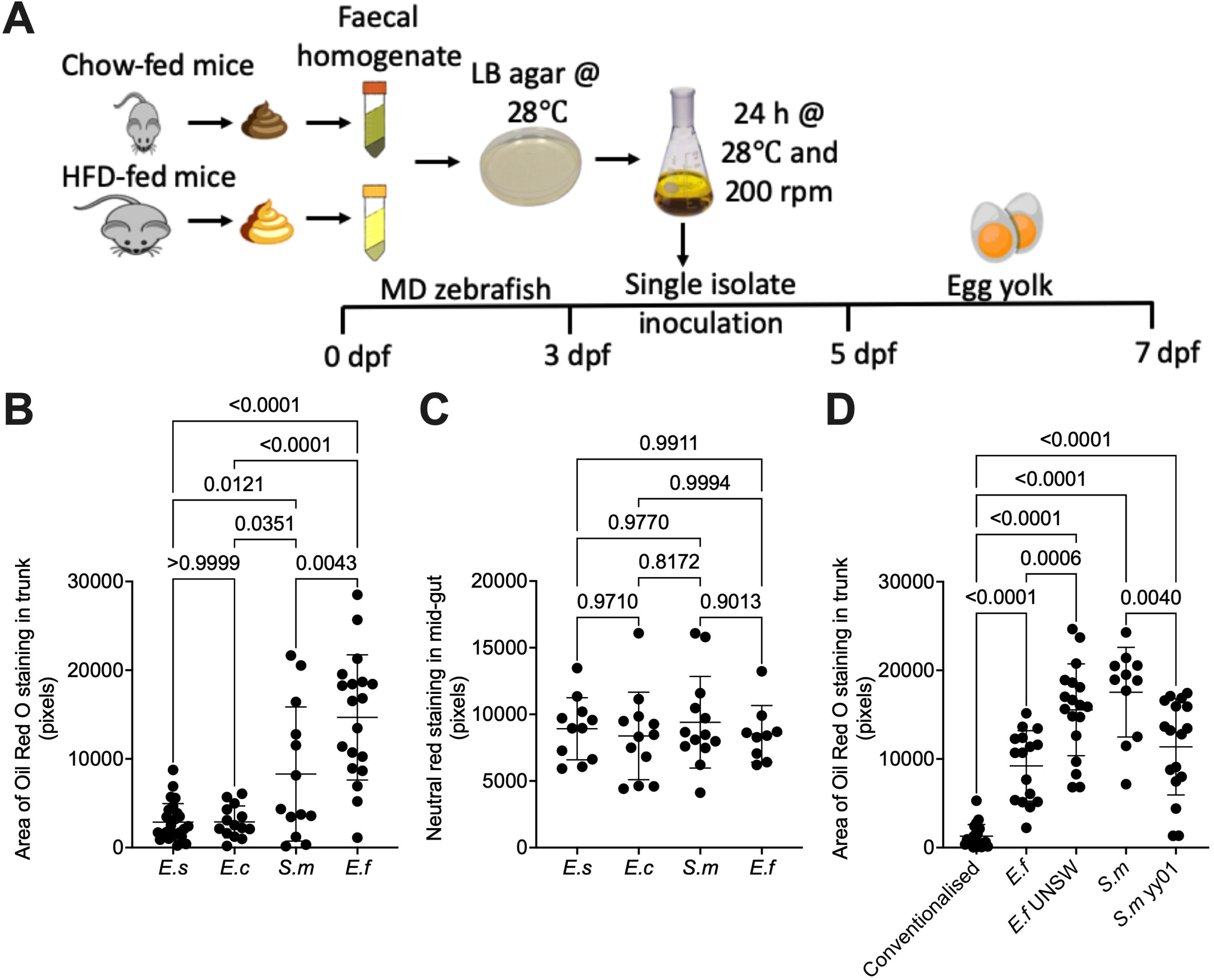
Identification of individual microbes with pathobiont activity. (a) Schematic describing workflow to isolate *Escherichia species* PYCC8248 (*E.s*), *Escherichia coli* strain Y15 (*E.c*), and *Enterococcus faecalis* strain YN771 (*E.f*) and inoculate into MD zebrafish with subsequent chicken egg yolk diet challenge. (b) Quantification of trunk vascular Oil Red O staining from the tail region posterior to the swim bladder in MD zebrafish inoculated with bacterial isolates *E.s*, *E.c*, and *E.f* and challenged with a chicken egg yolk diet from 5-6 dpf. (c) Area of neutral red stained in the mid-gut of 5 dpf gnotobiotic zebrafish embryos mono-associated with bacterial isolates *E.s*, *E.c*, and *E.f*. (d) Quantification of trunk vascular Oil Red O staining from the tail region posterior to the swim bladder in MD zebrafish inoculated with bacterial isolates *E.f* and *E. faecalis* UNSW 054400 type strain (*E.f* UNSW) then challenged with a chicken egg yolk diet from 5-6 dpf. Results are expressed as mean ± SD.

We colonised MD zebrafish larvae with individual isolates to determine if monoassociation could replicate the effects of bulk microbiota transfer. To determine if exposure of larvae to preparations of *E. faecalis, S. maltophilia*, or either of the *E. coli* strains resulted in gut colonisation we dissected guts from rinsed 5 dpf larvae after 2 days of monoassociation exposure and recovered bacteria onto LB agar. All tested strains yielded recoverable colonisation levels of approximately 100 CFU per larval gut. Gnotobiotic zebrafish larvae colonised with *E. faecalis* or *S. maltophilia* had increased Oil Red O staining compared to larvae colonised with either of the *Escherichia* strains after chicken egg yolk challenge (Figure 2B).

Interestingly, colonisation with *E. faecalis* or *S. maltophilia* did not increase the absorptive activity of the midgut intestinal epithelium compared to larvae colonised with either of the Escherichia strains suggesting the absorptive phenotype seen in bulk microbiota transplant is the product of multiple microorganisms or metabolites present in the complex mouse faecal microbiota preparations (Figure 2C).

We next sought to confirm our observations using a second strain of each bacterial species, *E. faecalis* UNSW 054400 type strain (*E.f* UNSW) and *S. maltophilia* yy01 (*S.m* yy01) isolated from another mouse in the same facility. Colonisation of MD zebrafish embryos with either strain replicated the hyperlipidaemic phenotype seen with our original isolates (Figure 2D). Interestingly, the type UNSW 054400 strain increased hyperlipidaemia beyond that seen with our YN771 isolate strain, while conversely the *S. maltophilia* yy01 strain was not as potent as our original CD103 strain, demonstrating strain-specific variability in our zebrafish embryo system.

### Stenotrophomonas maltophilia can accelerate hyperlipidaemia by digesting food external to the host

To identify the mechanisms by which *E. faecalis* and *S. maltophilia* were accelerating HFD-induced hyperlipidaemia in zebrafish larvae, we sought to investigate if the bacteria needed to be alive and/or in contact with the host.

We adapted our gnotobiotic methodology to expose MD embryos to heat killed bacterial preparations prior to chicken egg yolk feeding (Figure 3A). Exposure to heat-killed *E. faecalis* replicated the live *E. faecalis* hyperlipidaemic phenotype when compared MD embryos that had been exposed to either heat killed *S. maltophilia* or either of the *Escherichia* strains (Figure 3B). Conversely, heat-killed *S. maltophilia* did not induce hyperlipidaemia in concert with chicken egg yolk challenge.

**Figure 3:**
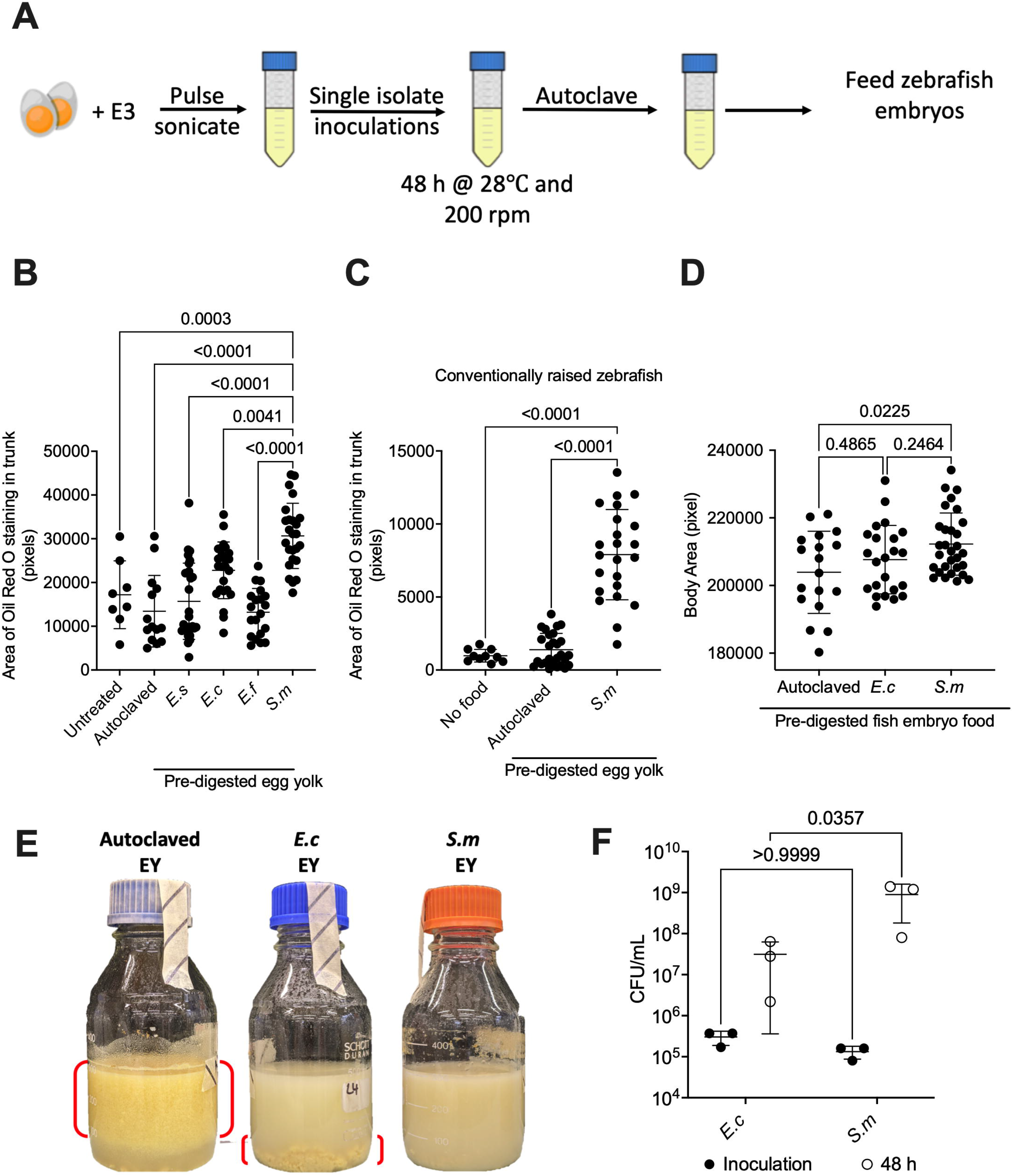
Stenotrophomonas maltophilia accelerates hyperlipidaemia by digesting food. (a) Schematic describing creation of “pre-digested” chicken egg yolk by incubation with bacterial isolates *E.s*, *E.c*, *S.m*, and *E.f*, autoclaving, for feeding to 5-7 dpf zebrafish embryos. (b) Quantification of trunk vascular Oil Red O staining from the tail region posterior to the swim bladder in MD zebrafish fed “pre-digested” egg yolk with bacterial isolates *E.s*, *E.c*, *S.m*, and *E.f* from 5-7 dpf. (c) Quantification of trunk vascular Oil Red O staining from the tail region posterior to the swim bladder in conventionally raised zebrafish fed “pre-digested” egg yolk with bacterial isolate *S.m* from 5-7 dpf. (d) Quantification of trunk vascular Oil Red O staining from the tail region posterior to the swim bladder in conventionally raised zebrafish fed “pre-digested” fish embryo food with bacterial isolates *E.c* and *S.m* from 5-7 dpf. (e) Representative images of “pre-digested” egg yolk with bacterial isolates *E.c* and *S.m* red brackets indicate fraction of the water column containing large particulates after autoclaving. (f) CFU recovery from chicken egg yolk “pre-digestion” reactions. Results are expressed as mean ± SD.

We hypothesised *S. maltophilia* might interact with the chicken egg yolk independently of the host colonisation. To test this hypothesis, we incubated chicken egg yolk with each of our 4 bacterial isolates in conditions representative of the zebrafish embryo media and then sterilised the “pre-digested” chicken egg yolk by autoclaving prior to feeding to MD zebrafish embryos (Figure 3C). Compared to either untreated chicken egg yolk, autoclaved chicken egg yolk, or chicken egg yolk incubated with the other three bacterial isolates, the chicken egg yolk that had been incubated with *S. maltophilia* increased hyperlipidaemia in zebrafish embryos (Figure 3D). A weaker, but statistically significant, effect was also seen with chicken egg yolk that had been incubated with *Escherichia coli* strain Y15-3 (*E.c*).

We next repeated this experiment in conventionally raised embryos and found the increased hyperlipidaemic potential of *S. maltophilia* “pre-digested” chicken egg yolk was replicated in larvae with a conventional microbiome (Figure 3E).

To examine the substrate specificity of *S. maltophilia*, we incubated commercially available fish embryo food with *Escherichia coli* strain Y15-3 or *S. maltophilia* and observed an increase in body size of only in embryos fed the commercial feed that had been “pre-digested” with *S. maltophilia* compared to untreated commercial feed (Figure 3F).

Visual inspection of *S. maltophilia* “pre-digested” chicken egg yolk suspensions suggested *S. maltophilia* had broken apart the chicken egg yolk resulting in smaller particles that could be more easily ingested and altered the biochemical properties of the chicken egg yolk as the solution contained much finer particles than for other treatments (Figure 3G). An intermediate phenotype was seen in chicken egg yolk that had been “pre-digested” by *E. coli* strain Y15-3. CFU recovery assays demonstrated higher growth of *S. maltophilia* than *E. coli* strain Y15-3 suggesting the better growth of *S. maltophilia* may convert chicken egg yolk into components that could be digested by zebrafish embryos (Figure 3H). We performed nutritional panel and free fatty acid analyses of chicken egg yolk that had been “pre-digested” by *S. maltophilia* or *E. coli* strain Y15-3 as an additional control (Table 1). These analyses revealed only a modest increase in energy content by *E coli* strain Y15-3 and *S. maltophilia*, and an increase in total fat content made up of monounsaturated and saturated fatty acids in *S. maltophilia*-incubated samples that was not seen in control or *E. coli* strain Y15-3-incubated samples.

These data illustrate a colonisation-independent mechanism by which *S. maltophilia* may enhance lipid uptake in zebrafish larvae by modifying food in the aqueous environment external to the host.

### Enterococcus faecalis accelerates hyperlipidaemia via host MyD88-mediated signalling

Our analyses had shown *E. faecalis* did not need to be alive but did need to be pre-associated with the hatching zebrafish larvae to accelerate hyperlipidaemia suggesting colonisation of the gut or other mucosal surfaces and subsequent recognition by the host may been necessary to accelerate hyperlipidaemia. Host innate immune signalling via the Myd88 adaptor protein is essential for the zebrafish intestinal epithelium to respond to microbial colonisation [14].

We performed knockdown of host *myd88* expression using multiple CRISPR-Cas9 gRNAs (Figure 4A). Host *myd88* expression was necessary for transducing the *E. faecalis*-induced hyperlipidaemic signal as *myd88* crispants had significantly less vascular Oil Red O staining than scrambled gRNA/Cas9-injected control embryos after colonisation with *E. faecalis* and chicken egg yolk challenge (Figure 4B).

**Figure 4:**
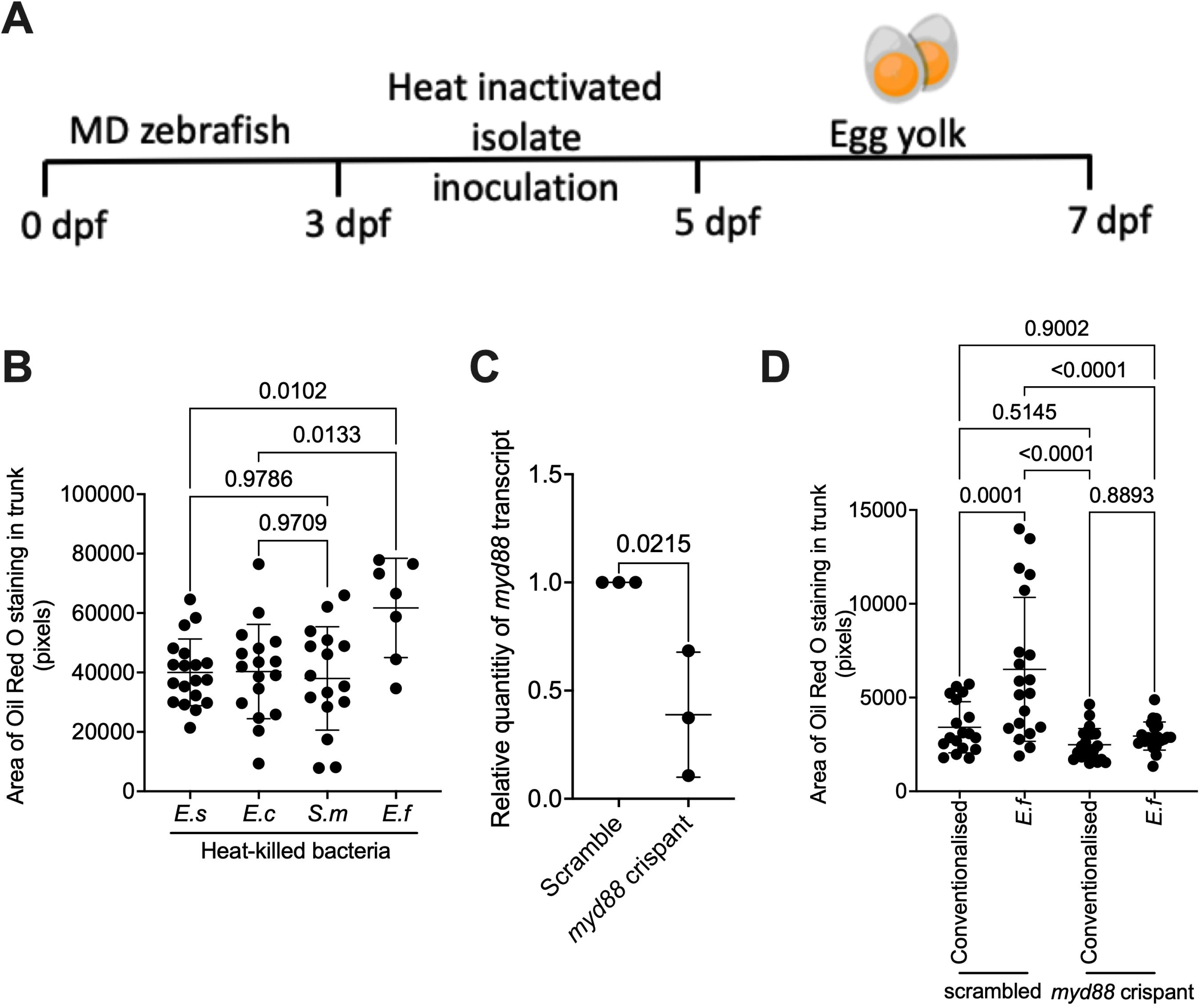
Enterococcus faecalis (E.f) accelerates hyperlipidaemia via host MyD88-mediated signalling. (a) Schematic describing inoculation of MD zebrafish embryos with heat-killed bacterial isolates *E.s*, *E.c*, and *E.f* from 3-5 dpf, followed by chicken egg yolk diet challenge from 5-7 dpf. (b) Quantification of trunk vascular Oil Red O staining from the tail region posterior to the swim bladder in MD zebrafish inoculated with heat killed bacterial isolates *E.s*, *E.c*, and *E.f* and challenged with a chicken egg yolk diet. (c) Quantification of *myd88* expression in zebrafish embryos injected with *myd88*-targeting CRISPR-Cas9 complexes at 5 dpf. Each dot represents a biological replicate of at least 10 embryos. (d) Quantification of trunk vascular Oil Red O staining from the tail region posterior to the swim bladder in control scrambled and *myd88* crispant embryos exposed to *E.f* and challenged with a chicken egg yolk diet. Results are expressed as mean ± SD.

### Gram positive cell wall components accelerate hyperlipidaemia in zebrafish embryos

*E. faecalis* is a Gram-positive bacterium. To determine if *E. faecalis*-driven hyperlipidaemia was due to a conserved Gram-positive cell wall component, we initially compared the hyperlipidaemic potential of heat-killed *E. faecalis* to heat killed *Staphylococcus xylosus* (*S.x)*, another Gram-positive bacterium obtained from the faecal microbiota of a mouse from the same facility. We found exposure of zebrafish larvae to heat-killed *S. xylosus* replicated the hyperlipidaemic effect of heat-killed *E. faecalis* after chicken egg yolk challenge (Figure 5A).

**Figure 5:**
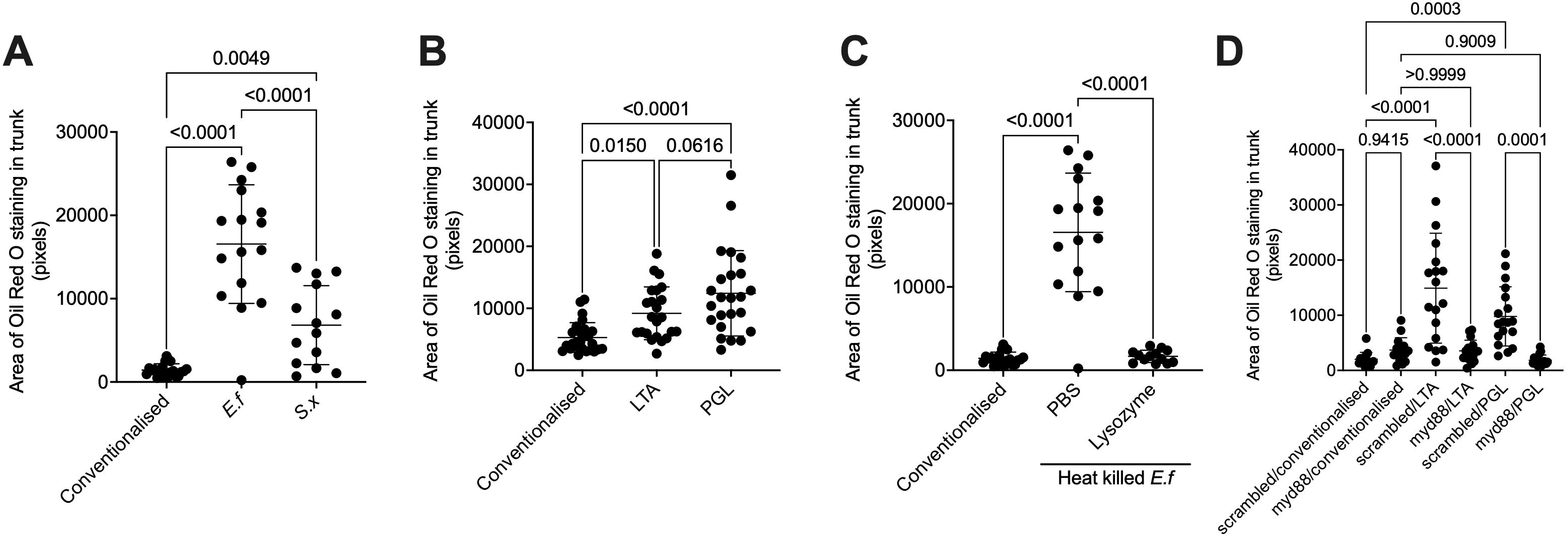
Gram-positive cell wall components accelerate hyperlipidaemia in zebrafish embryos. (a) Quantification of trunk vascular Oil Red O staining from the tail region posterior to the swim bladder in MD zebrafish inoculated with heat killed bacterial isolates *E.f* and Staphylococcus xylosus (*S.x*) from 3-5 dpf and challenged with a chicken egg yolk diet from 5-7 dpf. (b) Quantification of trunk vascular Oil Red O staining from the tail region posterior to the swim bladder in MD zebrafish pre-incubated with LTA or peptidoglycan from 3-5 dpf before co-challenge with chicken egg yolk diet from 5-7 dpf. (c) Quantification of trunk vascular Oil Red O staining from the tail region posterior to the swim bladder in MD zebrafish pre-incubated with lysozyme-treated heat killed *E.f* from 3-5 dpf and challenged with a chicken egg yolk diet from 5-7 dpf. (d) Quantification of trunk vascular Oil Red O staining from the tail region posterior to the swim bladder in control scrambled and *myd88* crispant embryos pre-incubated with LTA or peptidoglycan from 3-5 dpf before co-challenge with chicken egg yolk diet from 5-7 dpf. Results are expressed as mean ± SD.

Next we directly soaked MD larvae in a sublethal dose of 25 μg/mL purified lipoteichoic acid (LTA) or peptidoglycan (PGL), which are major components of the Gram-positive cell wall, prior to challenge with chicken egg yolk feeding [15]. Either one of these purified ligands were able to accelerate hyperlipidaemia in MD larvae (Figure 5B).

To test the requirement for intact *E. faecalis* PGL to accelerate hyperlipidaemia, we exposed MD larvae to lysozyme-digested heat-killed *E. faecalis* prior to chicken egg yolk feeding. Lysozyme digestion ablated the ability of heat-killed *E. faecalis* to accelerate hyperlipidaemia (Figure 5C).

To determine if host innate immune signalling via the Myd88 adaptor protein transduced the LTA or PGL-induced signal that accelerates hyperlipidaemia, we again knocked down *myd88* using a pooled CRISPR-Cas9 gRNA approach and exposed crispants to LTA or PGL. Depletion of host *myd88* ablated the sensitivity of embryos to LTA and PGL-induced accelerated hyperlipidaemia (Figure 5D).

Together, these experiments demonstrate larval zebrafish lipid metabolism is sensitive to the presence of Gram-positive bacterial cell wall components via Myd88-mediated host signalling pathways.

## Discussion

Our study demonstrates the utility of the gnotobiotic zebrafish platform to screen donor microbiota samples for transplantable biological activities in combination with an exogenous environmental factor. The addition of an exogenous trigger to the experimental system is an important permutation that allows the identification of microbes whose interactions with the host only become apparent in combination with an environmental challenge such as diet.

We applied the gnotobiotic zebrafish platform to explore the diet-hyperlipidaemia axis as there is a growing interest in microbiome studies within the cardiovascular disease field [16]. The gnotobiotic zebrafish platform represents a disruptive technology that could be adapted to identify pathobiont species from human cardiovascular disease patients and as a “first pass” platform in mechanistic studies with appropriate zebrafish models of cardiovascular pathology.

Our investigation of *E. faecalis*-accelerated hyperlipidaemia uncovered a surprising role of Gram-positive cell wall component-triggered Myd88 signalling in zebrafish embryo lipid metabolism. The presence of this response in zebrafish embryos has broad implications for the use of zebrafish embryos to study transplanted mammalian microbiota as it may not be representative of the mammalian response to colonisation with Gram positive organisms. Most zebrafish embryo mono-association studies have been carried out with Gram negative organisms including *E. coli*, *A. veronii*, *V. cholerae*, and *P. aeruginosa* [17–19].

The comparison of heat killed *E. faecalis* to heat killed *S. xylosus* demonstrated an increased ability of heat killed *E. faecalis* to accelerate diet-induced hyperlipidaemia. This suggests that there is variability in hyperlipidaemia accelerating potential amongst Gram positive organisms in zebrafish embryos. The basis of this difference could be further explored using model Gram positive organisms and Gram-positive organisms that are natural commensals in the zebrafish gut. A recent paper by Griffin *et al*. has demonstrated a role for *E. faecalis* SagA enzyme in producing immuno-stimulatory muropeptides [20], as we have previously demonstrated a conserved function of the zebrafish NOD orthologs it is possible that SagA-mediated production of muropeptides from bacterial peptidoglycan contributes to our hyperlipidaemia-accelerating phenotype.

*Caenorhabditis elegans* fed with *S. maltophilia* accumulate neutral lipids within intracellular lipid droplets driven by a bacterially-encoded mechanism that is independent of the innate immune response [21]. Our finding that two distinct mouse-associated *S. maltophilia* strains were able to increase the hyperlipidaemic potential of chicken egg yolk suggests the digestive ability of *S. maltophilia* is potentially conserved between strains and potentially between host species. Previous studies have identified *S. maltophilia* strains within zebrafish gut microbiomes [9], these zebrafish-associated strains could be analysed to determine if increased lipid uptake is a consequence of natural host-*S. maltophilia* pairs and for comparative studies to identify the food modifying mechanisms employed by our *S. maltophilia* CD103 strain isolate.

We also found an embryo growth-enhancing effect of pre-digesting commercially available zebrafish embryo feed with *S. maltophilia* which suggests *S. maltophilia* could have commercial applications as an aquaculture feed additive. The use of *S. maltophilia* as a feed additive is potentially risky as this organism is associated with opportunistic infections in humans and is resistant to a wide range of antibiotics so care should be taken to avoid cross over into the food chain [22, 23].

As a proof-of-principle study, our work demonstrates the feasibility of studying the interaction of bacterial species transplanted from a mammalian host with an environmental factor to result in a conditional phenotype in the tractable zebrafish model system. Further work is required to examine effects of the two pathobiont species identified in our study on mammalian models of hyperlipidaemia and to correlate their colonisation with the progression of cardiovascular disease phenotypes in mammalian models and human samples.

## Methods

### Zebrafish handling

Adult zebrafish were housed at the Centenary Institute (Sydney Local Health District Animal Welfare Committee Approval 17-036). Zebrafish embryos were obtained by natural spawning and conventionally raised embryos were maintained in E3 media at 28°C.

### Generation of microbiome-depleted (MD) zebrafish embryos

Microbiome-depleted (MD) zebrafish were created and maintained as previously described [24]. Briefly, freshly laid embryos were rinsed with 0.003% v/v bleach in sterile E3 and rinsed 3 times with sterile E3. Bleached embryos were raised in sterile E3 supplemented with 50 μg/mL ampicillin (Sigma), 5 μg/mL kanamycin (Sigma) and 250 ng/mL amphotericin B (Sigma) in sterile tissue culture flasks. Dead embryos and chorions were aseptically removed at one and three dpf respectively.

Conventionalised zebrafish were used as a control, where 3 dpf MD zebrafish were inoculated with system water from the aquarium at Centenary Institute.

### Generation of mouse faecal microbiota specimens

Mice were housed at the Centenary Institute (Sydney Local Health District Animal Welfare Committee Approval 2018/016). C57BL/6J mice were housed in a pathogen-free and temperature-controlled environment, with 12 hours of light and 12 hours of darkness, and free access to food and water. Mice were provided with a High Fat Diet (HFD) or chow from 6 to 30 weeks of age. The HFD was prepared in-house based on rodent diet no. D12451 (Research Diets New Brunswick, USA) and its calories were supplied as: fat 45%, protein 20%, and carbohydrate 35% [25]. The chow diet was commercially produced by Speciality Feeds as “Irradiated Rat and Mouse Diet” and its calories were supplied as: fat 12%, protein 23%, and carbohydrate 65%.

Faecal pellets were collected from mice that were housed in different cages. Individual faecal pellets were collected into sterile 1.7 mL microcentrifuge tubes and homogenised in 1 mL of sterile E3 by pipetting. Homogenised specimens were centrifuged at 500 G for 2 minutes to sediment fibrous material and the supernatant was collected. Supernatants were supplemented with glycerol to a final concentration of 25% v/v, aliquoted, and frozen at −80°C for experimental use.

### Colonisation of zebrafish with mouse faecal microbiota

MD zebrafish were colonised at 3 dpf by transfer into sterile E3 and addition of 200 μL thawed faecal homogenate supernatant. At 5 dpf, embryos were rinsed with E3 and placed on a chicken egg yolk diet.

### Neutral red staining and morphology measurements

Neutral red staining was performed as previously described [26]. Briefly, 2.5⍰μg/mL neutral red was added to the media of 4 dpf embryos and incubated overnight. Embryos were rinsed with fresh E3 to remove unbound neutral red and live imaged on a Leica M205FA microscope with a consistent zoom between specimens in a single experiment. The area of neutral red-stained midgut was measured in pixels using ImageJ. Body size was measured in pixels using ImageJ.

### Zebrafish high fat diet challenge with chicken egg yolk

High fat diet challenge with chicken egg yolk was performed as previously described [13]. Briefly, 5 dpf zebrafish embryos were placed in an E3 solution containing 0.05% w/v emulsified hard boiled chicken egg yolk in glass beakers. Beakers were housed in a 28°C incubator with a 14:10 hour light:dark cycle. The emulsified hard boiled chicken egg yolk solution was changed daily.

### Oil Red O staining assay

Oil Red O staining and analysis was performed as previously described [13, 27, 28]. Briefly, 7 dpf embryos were fixed overnight at 4°C in 4% paraformaldehyde, rinsed with PBS, and rinsed stepwise through to propylene glycol. Embryos were stained with filtered 0.5% (w/v) Oil Red O dissolved in propylene glycol overnight at room temperature. Unbound dye was removed by washing with propylene glycol and embryos were rinsed stepwise through to PBS for imaging.

Embryos were imaged on a Leica M205FA microscope. Experimental batches of colour images were analysed in ImageJ by adjusting the colour threshold function to eliminate non-red signal, this output was then converted to a binary mask and the tail region posterior to the swim bladder was selected to measure the area of particles.

### Isolation, identification, and handling of bacterial isolates from mouse faecal microbiota samples

Faecal homogenate supernatants were plated onto LB Agar (Amyl Media) and incubated at 28°C for two days. Individual isolates were grown in broth culture in Luria Broth (Miller’s LB Broth Base, Thermofisher) at 28°C overnight with 200 RPM shaking.

For identification, bacteria were harvested from broth culture and subjected to PCR using universal 16S primers (5⍰-3⍰) Fw Rv. PCR products were sequenced by Sanger Sequencing (AGRF) and NCBI records were searched by BLAST to identify the closest matching bacterial strain by sequence identity.

For gnotobiotic experiments, bacteria were harvested from overnight broth cultures and resuspended in sterile E3 zebrafish embryo media at a concentration of OD600 0.2. MD zebrafish in autoclaved E3 were then inoculated with the bacterial suspension at a ratio of 1:200.

For heat killed bacterial inoculations, bacteria were resuspended in E3 at a concentration of OD600 0.2 and heat-killed in a 95°C heat block for 30 minutes. Heat killed bacterial solutions were then added at 1:200 ratio to 3 dpf MD zebrafish.

For purified ligand exposure, 3 dpf MD zebrafish were soaked in 25 μg/mL lipoteichoic acid (Sigma) or 25 μg/mL peptidoglycan (Sigma).

For lysozyme digestion of heat-killed *E. faecalis*, a suspension of heat-killed bacteria was incubated with 50 μg/mL lysozyme (Sigma) at 37°C for 6 h before being used to inoculate 3 dpf MD zebrafish at a ratio of 1:200.

### Digestion of chicken egg yolk

20 g hard-boiled chicken egg yolk was mixed with 40 mL E3 and emulsified using a Branson Digital Sonifier sonicator. Emulsified egg yolk was inoculated with bacterial isolates in E3 (described above) at a 1:200 ratio. Egg yolk mixture was then placed in a shaker at 28 °C at 200 rpm and for 48 h. Samples were then autoclaved prior to feeding experiments.

### Analysis of chicken egg yolk composition

“Pre-digested” samples were autoclaved and mailed to Australian Laboratory Services (VIC, Australia) for commercial grade nutritional analyses.

### Gene knockdown with CRISPR-Cas9

gRNA templates for *myd88* (5’-3’): Target 1 TAATACGACTCACTATAGGCAGTTTCCGAAAGAAACTGTTTTAGAGCTAGAAATAGC, Target 2 TAATACGACTCACTATAGGAAAAGGTCTTGACGGACTGTTTTAGAGCTAGAAATAGC, Target 3 TAATACGACTCACTATAGGAACTGTTTGATCATCTCGGTTTTAGAGCTAGAAATAGC, Target 4 TAATACGACTCACTATAGGTTTTTTCGATAAGCTCACGTTTTAGAGCTAGAAATAGC. gRNA was synthesized as previously described [29].

A 1:1 solution of gRNA and 500 μg/mL of Cas9 nuclease V3 (Integrated DNA Technology, Sigma, or or Sydney Analytical) was prepared with phenol red dye (Sigma, P0290). Freshly laid eggs were collected from breeding tanks and the solution was injected in the yolk sac of the egg before the emergence of the first cell with a FemtoJet 4i (Eppendorf).

Knockdown efficacy was monitored by RT-qPCR as previously described [30]. *myd88*-specific primers (5’-3’): Fw ACAGGGACTGACACCTGAGA, Rv GACGACAGGGATTAGCCGTT.

To derive MD crispant embryos, injected embryos were placed in E3 containing ampicillin, kanamycin and amphotericin B as described previously. Bleaching injected embryos caused high mortality rates.

### Statistics

All statistical analyses (t-tests and ANOVA where appropriate) were performed using GraphPad Prism 8. Outliers were removed using ROUT, with Q = 1%. All data shown are representative of at least 2 biological replicates.

## Data availability

Source data are provided with this paper.

Raw image and analysis data is archived for 10 years by The Centenary Institute and available on request from the corresponding author (Sydney, Australia).

## Acknowledgements

Funding: Australian National Health and Medical Research Council Project Grant APP1099912; The University of Sydney Fellowship G197581; NSW Ministry of Health under the NSW Health Early-Mid Career Fellowships Scheme H18/31086 to SHO. Australian National Health and Medical Research Council Centres of Research Excellence Grant APP1153493 to WJB. The funders had no role in study design, data collection and analysis, decision to publish, or preparation of the manuscript.

The *E. faecalis* UNSW 054400 type strain was provided by Dr Laurence Marcia.

The authors acknowledge the facilities and the technical assistance of Dr Angela Kurz at the BioImaging Facility and Sydney Cytometry at Centenary Institute, and Dr Mario Torrado Del Rey of Sydney Analytical at The University of Sydney for supply of some Cas9 protein.

The authors declare no conflicts of interest.

