## Supplementary material for "Transplantation of high fat fed mouse microbiota into zebrafish larvae identifies MyD88-dependent acceleration of hyperlipidaemia by Gram positive cell wall components": Table 1

Table 1: Nutritional content of pre-digested egg yolk after 48 hours incubation.

| Sample | Control | *E. coli* strain Y15-3 | *S. maltophilia* |
| --- | --- | --- | --- |
| Moisture (g/100 mL) | 98.4 | 98.1 | 98.2 |
| Ash (g/100 mL) | 0.1 | 0.1 | 0.1 |
| Carbohydrates (g/100 mL) | 0.2 | <0.1 | 0.2 |
| Total sugars (g/100 mL) | <0.1 | <0.1 | <0.1 |
| Energy (kJ/100 mL) | 44 | 49 | 51 |
| Total fat (g/100 mL) | 0.9 | 0.9 | 1.1 |
| Monounsaturated fatty acids (g/100 mL) | 0.5 | 0.5 | 0.6 |
| Polyunsaturated fatty acids (g/100 mL) | 0.1 | 0.1 | 0.1 |
| Saturated fatty acids (g/100 mL) | 0.3 | 0.3 | 0.4 |
| Trans fatty acids (g/100 mL) | <0.1 | <0.1 | <0.1 |
| Sodium (mg/100 mL) | 14 | 14 | 14 |
| Protein (Dumas, g/100 mL) | 0.4 | 0.9 | 0.4 |
| Free fatty acids (%) | <0.03 | <0.03 | <0.03 |
